# High-speed whole-genome sequencing of a Whippet: Rapid chromosome-level assembly and annotation of an extremely fast dog’s genome

**DOI:** 10.1101/2024.08.16.608262

**Authors:** Marcel Nebenführ, David Prochotta, Alexander Ben Hamadou, Axel Janke, Charlotte Gerheim, Christian Betz, Carola Greve, Hanno J. Bolz

**Author notes:** Authors contributed equally (Marcel Nebenführ, David Prochotta). Authors for correspondence: Marcel Nebenführ, David Prochotta, Hanno J. Bolz.

## Abstract

**Background:** The time required for sequencing and *de novo* assembly of genomes is highly dependent on the interaction between laboratory work, sequencing capacity, and the bioinformatics workflow. As a result, genome projects are often not only limited by financial, computational and sequencing platform resources, but also delayed by second party sequencing service providers. By bringing together academic biodiversity institutes and a medical diagnostics company with extensive sequencing capabilities and know-how, we aimed at generating a high-quality mammalian *de novo* genome in the shortest possible time period. Therefore, we streamlined all processes involved and chose a very fast dog as a model: The Whippet.

**Findings:** We present the first chromosome-level genome assembly of the Whippet. We used PacBio long-read HiFi sequencing and reference-guided scaffolding to generate a high-quality genome assembly. The final assembly has a contig N50 of 55 Mbp and a scaffold N50 of 65.7 Mbp. The total assembly length is 2.47 Gbp, of which 2.43 Gpb were scaffolded into 39 chromosome-length scaffolds. In addition, we used available mammalian genomes and transcriptome data to annotate the genome assembly. The annotation resulted in 28,383 transcripts resembling a total of 90.9% complete BUSCO genes and identified a repeat content of 36.5%.

**Conclusions:** Sequencing, assembling, and scaffolding the chromosome-level genome of the Whippet took less than a week and adds a high-quality reference genome to the list of domestic dog breeds sequenced to date.

## Data description

### Background information

Although recent advances in sequencing technologies have opened up the field of genomics for a broader scientific community, sequencing and assembling large eukaryotic genomes of non-model organisms remains challenging. Even with access to samples of sufficient quality, limited funding for sequencing, lack of computing power and/or sequencing technology and capacity and inevitable outsourcing can significantly increase the turnaround times.

In response to the accelerating global biodiversity loss, national and international initiatives have expanded genomic reference resources for biodiversity research and conservation. The use of reference genomes in population genomics facilitates the characterization of genetic diversity and adaptation through local variant enrichment, providing a basis for biodiversity assessment, conservation and restoration [1–3].

Furthermore, non-human genomes have become increasingly helpful for the interpretation of human genetic variants and their relevance for disease [4,5]; this particularly applies to *de novo*-assembled long-read human genomes that - in contrast to short-read genomes - allow for the identification of complex (structural) variants that would otherwise escape detection. In both biodiversity and conservation research as well as in medical genetics, the timely generation and assembly of genomic data is of utmost importance.

We joined forces between an academic biodiversity institute and a medical diagnostic company with extensive sequencing capacity and experience to generate a rapid chromosome-level *de novo* genome of the Whippet. We demonstrate that streamlined laboratory and bioinformatics workflows with PacBio HiFi long-read whole-genome sequencing (LR-WGS) enable the generation of a high-quality reference genome within one week (**Fig. 1**). Besides this proof-of-concept study of the rapid Whippet genome, our collaboration includes continuous LR-WGS of *de novo* genomes of various endangered species (including non-vertebrates and plants). Our study may serve as a paradigm for such cooperations applicable to a wide range of human and non-human genome projects, from biodiversity research to domestic animal and agricultural research.

**Fig. 1.**
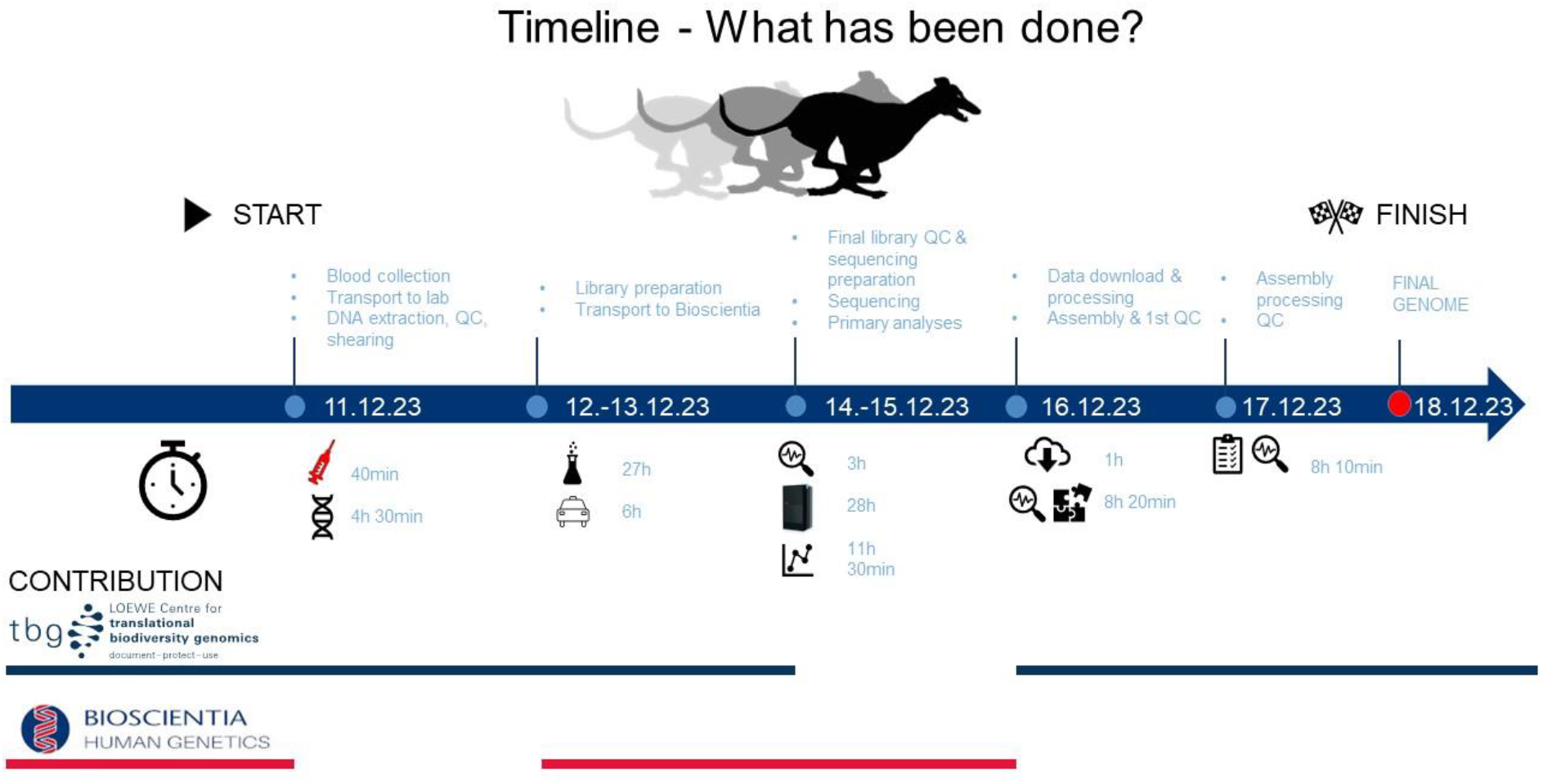
Project timeline with day-by-day progress description and time requirements. The blue line represents the contribution of the biodiversity research centre (TBG) at each step and the red line represents the contributions made by the medical diagnostics company (Bioscientia).

### Sampling, DNA extraction, and sequencing

High molecular weight genomic DNA was extracted from peripheral blood leukocytes of a three-year-old male dog (*Canis lupus familiaris*), a Whippet, using the PacBio Nanobind CBB kit (Pacific Biosciences, Menlo Park, CA). Blood was taken during a routine veterinary procedure, collected in EDTA-coated vials and frozen at −20°C. DNA concentration and DNA fragment lengths were evaluated using the Qubit dsDNA BR Assay kit on the Qubit Fluorometer (Thermo Fisher Scientific, Waltham, MA) and the Genomic DNA Screen Tape on the Agilent 4150 TapeStation system (Agilent Technologies, Santa Clara, CA), respectively. Two SMRTbell libraries were prepared according to the instructions in the SMRTbell Express Prep Kit v3.0. The final concentrations were 68 ng/μl and 76 ng/μl, with a total input of approximately 10 µg of sheared DNA per library. Annealing of sequencing primers, binding of sequencing polymerase, and purification of polymerase-bound SMRTbell complexes were performed using the Revio polymerase kit (PacBio, Menlo Park, CA, USA). Loading concentration for sequencing was 250 pM.

### Genome assembly and polishing

The two sequencing runs on a PacBio Revio^®^ instrument of the Whippet yielded a total of ∼128 Gbp of sequence data, with an average subread N50 of ∼17.8 kbp. We assembled the genome using Hifiasm v0.18.8-r525 [6] with default settings and scanned the initial genome assembly for contamination using FCS-GX v0.4.0 [7].

We polished the raw assembly three times using Inspector v1.0.1 [8] which includes Flye v2.9.3 [9] for structural error correction.

After polishing, we performed reference-guided scaffolding using RagTag v2.1.0 [10] with default settings. For that, the NCBI reference genome for dogs was chosen as reference, which is the German Shepherd dog genome (GCA_011100685.1). Afterwards, we used TGS-GapCloser v1.2.1 [11] to close assembly gaps using three rounds of gap-filling and the Racon v1.0.5 [12] module for polishing the assembly gaps.

### Assembly QC

We evaluated both the raw assembly and the final assembly with Merqury v1.3 [13] in combination with Meryl v1.4.1, as well as Inspector v1.0.1.

We calculated assembly contiguity statistics of the final genome using Inspector and performed a gene set completeness analysis of the Whippet genome and other available dog genomes for comparison using BUSCO v5.4.22 [14] with the provided Carnivora orthologous genes database (carnivora_odb10).

We mapped the reads back to the genome using minimap2 v2.28 [15] with the ‘-ax map-hifi’ option to output a SAM file. The SAM file was sorted and converted to a BAM file with samtools v1.20 [16]. We then marked duplicate sequences with sambamba v1.0.1 [17] and analyzed mapping statistics with QualiMap v2.2.1 [18].

The initial assembly, based only on the HiFi long reads, already recovered 36 (of 39) chromosome-sized scaffolds. This further demonstrates how effective and therefore invaluable accurate long-read sequencing has become in mammalian genomics.

The final polished, scaffolded, and gap-closed assembly consists of 148 scaffolds with a total length of 2.47 Gbp, while 2.43 Gbp were placed into 39 chromosome-sized scaffolds (**Fig. 2A,B**; **Tab. 1**). Gene completeness analysis of the genome based on BUSCO’s Carnivora dataset identified 14,170 complete single-copy orthologous sequences, corresponding to 97.72 % completeness, and 77 (0.53 %) missing genes (**Fig. 2A**).

**Fig. 2.**
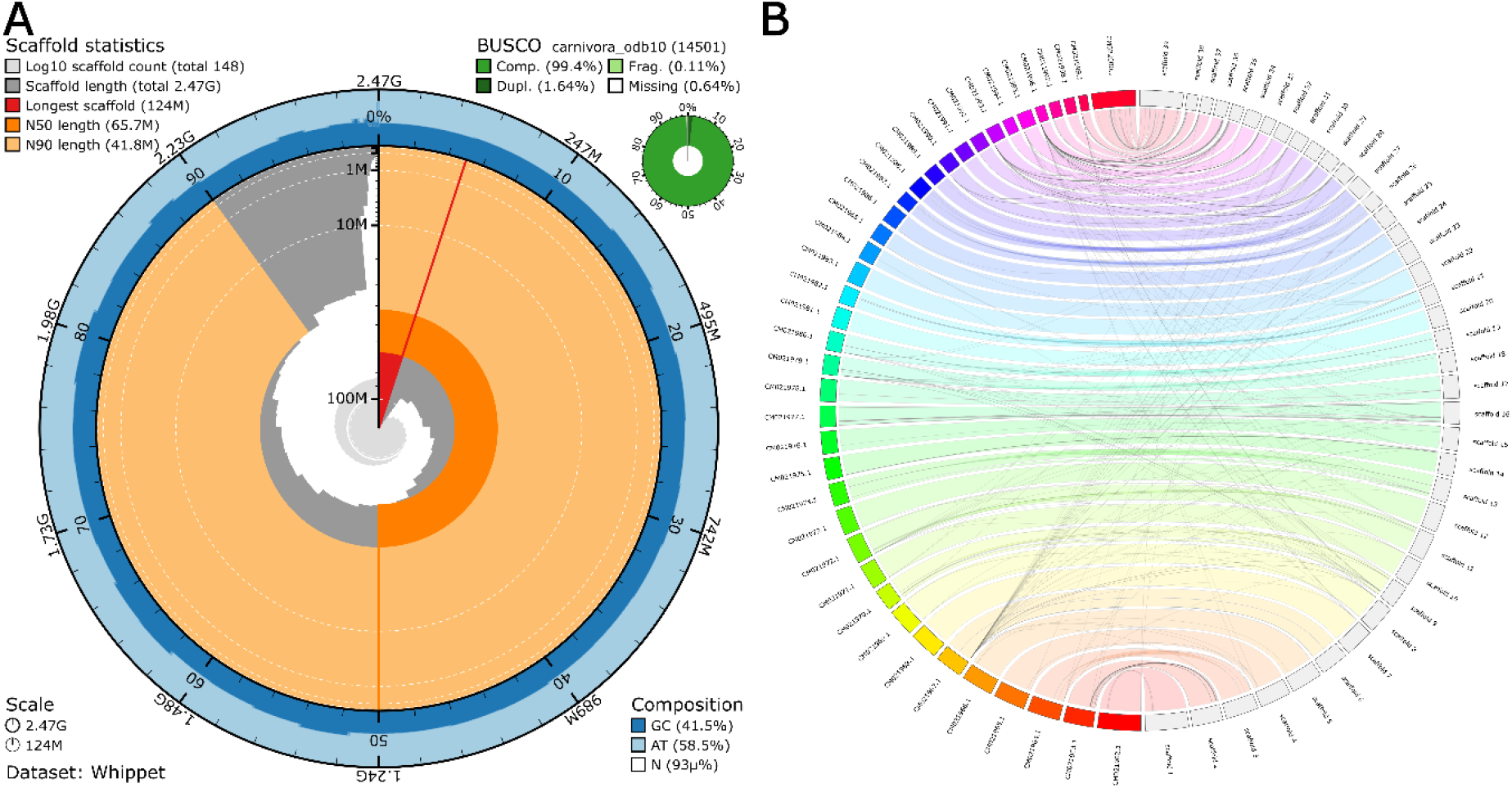
**(A)**. Snailplot based on the polished and reference-scaffolded assembly showing BUSCO gene completeness results and basic assembly statistics. **(B)**. Whole-genome synteny between the chromosome-level assembly of the German Shepherd and our chromosome-level assembly of the Whippet.

**Tab. 1.**
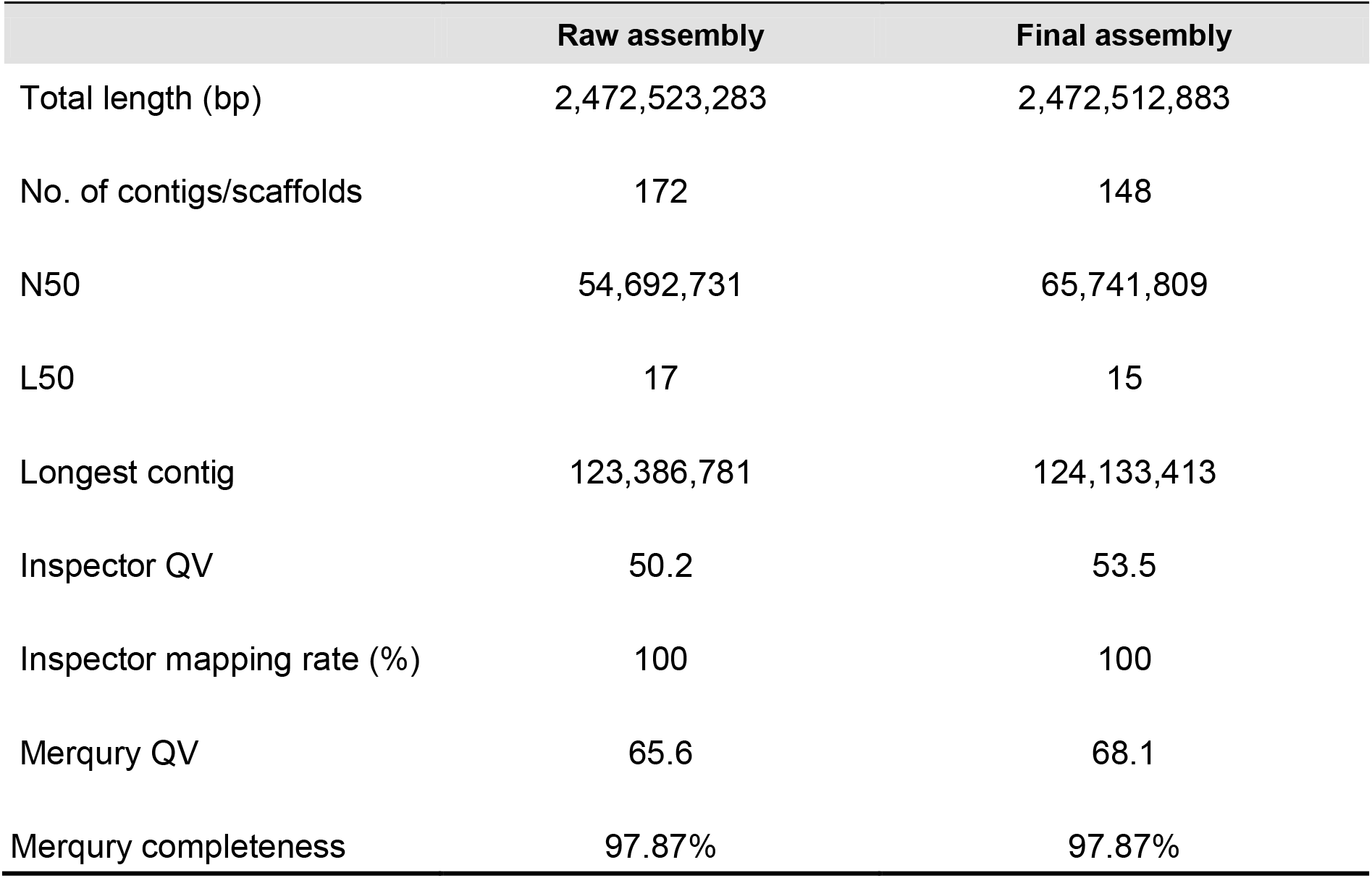
Basic assembly statistics, Inspector and Merqury quality values of the raw and final Whippet assembly.

|  | Raw assembly | Final assembly |
| --- | --- | --- |
| Total length (bp) | 2,472,523,283 | 2,472,512,883 |
| No. of contigs/scaffolds | 172 | 148 |
| N50 | 54,692,731 | 65,741,809 |
| L50 | 17 | 15 |
| Longest contig | 123,386,781 | 124,133,413 |
| Inspector QV | 50.2 | 53.5 |
| Inspector mapping rate (%) | 100 | 100 |
| Merqury QV | 65.6 | 68.1 |
| Merqury completeness | 97.87% | 97.87% |

Both Inspector and Merqury, coupled with the high BUSCO score, indicate a highly contiguous, accurate and complete genome. In addition, available dog assemblies were downloaded to compare the BUSCO completeness across published reference genomes (**Fig. 3, Tab. S2**). In addition, the qualimap report for the resulting BAM file showed 99.98% of mapped reads (8,332,294), a mean mapping quality of 49.7, a mean coverage of 55.1X, and an error rate of 0.0057.

**Fig. 3.**
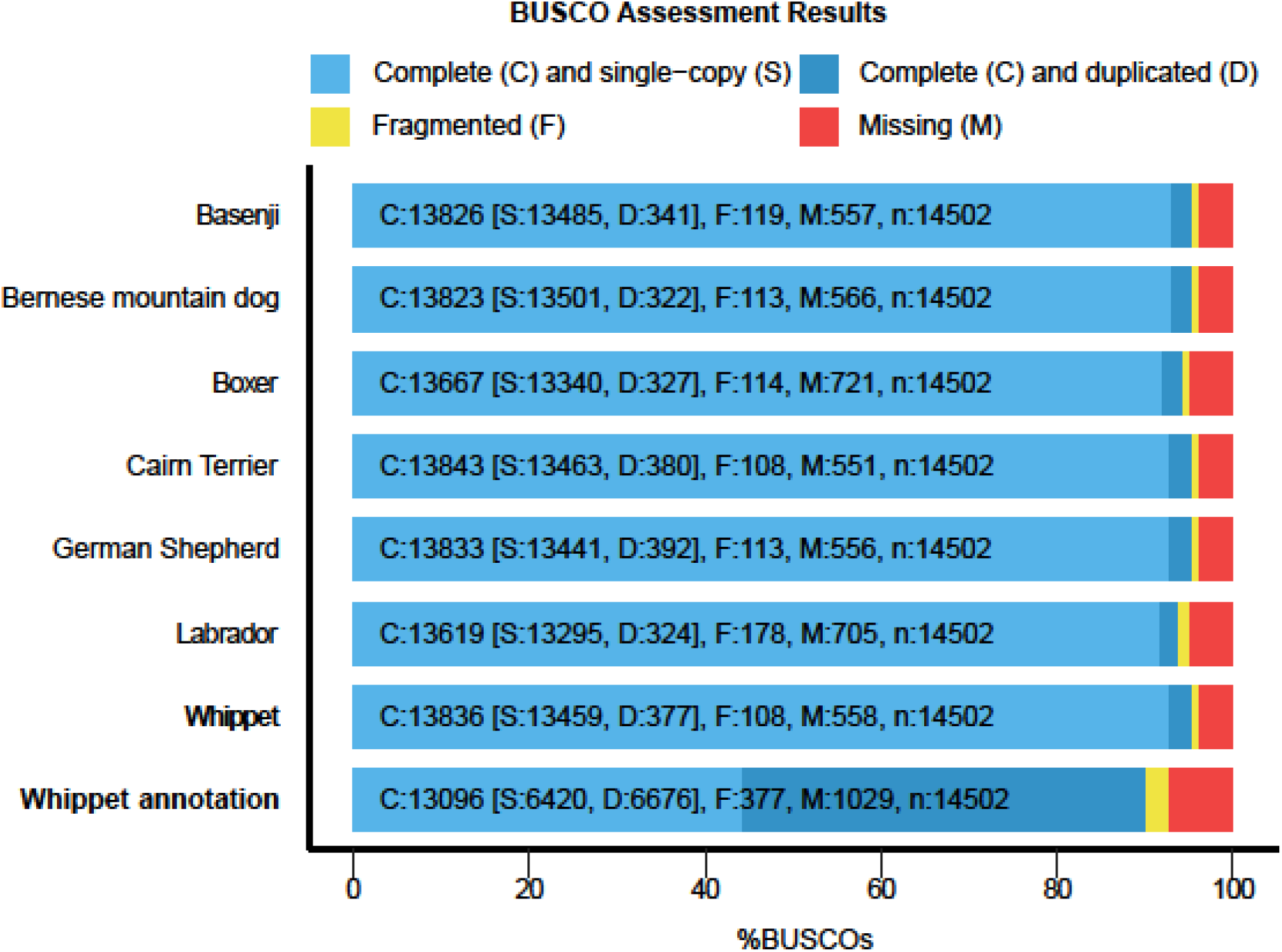
Comparison of BUSCO completeness statistics based on the carnivora database between our final Whippet assembly and annotation, and other available dog assemblies. Complete single-copy genes are shaded light blue, complete duplicated sequences are shaded blue. Fragmented genes are shaded yellow, and missing sequences are shaded red. The numbers of complete single copy (S), complete duplicated (D), fragmented (F) and missing genes (M) for the respective genome are shown in each column. The total number of genes in the BUSCO carnivora library is denoted as n.

### Heterozygosity

To calculate genome wide heterozygosity in the Whippet genome, we first mapped the reads used for assembly to the genome and marked duplicates using sambamba v1.0.0, then counted base depth at all sites using sambamba v1.0.0. We then estimated the Site Frequency Spectrum (SFS) with ANGSD v0.940 [19] and used the output files to run realSFS with 200 bootstrap replicates to calculate the folded SFS. From this we calculate the heterozygosity by dividing the heterozygous sites by the sum of the homozygous and heterozygous sites. In line with other dog genomes the heterozygosity is at 0.09%. Our heterozygosity analysis delivers standard baseline data for a *de novo* sequenced genome. It allows for a first glimpse into the genetic diversity of Whippets. However, it cannot be representative for Whippets in general.

## Genome annotation

### Repeat annotation

To annotate repetitive regions in the Whippet genome, we used RepeatModeler v2.1 [20] to create a *de novo* repeat library for our assembly. Afterwards, we used RepeatMasker v4.1.6 [21] to hardmask repeats based on the modeled repeats. Our analyses identified 36.5% of repeats in the genome, of which the majority consisted of LINEs (20.32%) and LTR elements (3.68%). In addition, 5.37% of unclassified elements were identified (**Tab. S2**).

### Gene annotation

Gene annotation was performed on the unmasked assembly using GeMoMa v1.9 [22]. Therefore, available annotated high-quality genomes of dogs and other mammals were used as references to identify genes (**Tab. S1**). The final gene annotation resulted in 28,383 transcripts. In addition, BUSCO analysis identified 90.9% complete BUSCOs, suggesting a high annotation completeness (**Fig. 3**).

### Myostatin

Since a founder mutation in myostatin (*MSTN*) has been reported in Whippet dogs [23], we analyzed our Whippet genome for this mutation by comparing its *MSTN* sequences to the wild-type references of *Canis lupus familiaris* (AY367768) and found no mutation in the sequence (**Fig. S3**). We first identified the position of the gene within the genome with MMseqs2 [24] and then visualized the region in IGV [25] to check for mutations in the aligned reads. That said, the analysis of our Whippet genome‘s *MSTN* sequence shall primarily exemplify a possible application of available high-quality genomes, especially for breeding purposes, but is not representative for Whippets in general.

## Conclusion

In our proof-of-concept study, we show that teaming up of a medical diagnostics company with a biodiversity research institute may deliver extremely rapid *de novo*-assembled HiFi long-read genomes. This was possible through close and streamlined time management and collaboration including all required participants for a genome project, namely a veterinarian, laboratory facilities, the sequencing facility, and the bioinformatics unit.

## Data availability

All raw data generated in this study are accessible at GenBank under *BioProject* PRJNA1114051. Annotation, results files and other data are available in the GigaDB repository.

## Author contributions

A.B.H. and Ch.G. performed the DNA extraction and the library preparation. D.P. and M.N. assembled and analyzed the genomes, and conducted the downstream analyses. C.G., M.N., D.P. and H.J.B. jointly supervised the project and wrote the manuscript with input from A.B.H., A.J., C.G., and C.B. All authors read and approved the final manuscript before submission.

## List of abbreviations

BUSCO: Benchmarking Universal Single-Copy Orthologs
CCS: Circular Consensus Sequencing
HiFi: High Fidelity
LINEs: Long interspersed nuclear elements
LR-WGS: Long-Read Whole-Genome Sequencing
LTR: Long Terminal Repeat
SFS: Site Frequency Spectrum

## Conflict of Interest

The authors declare that they have no competing interests.

## Consent for publication

Not applicable.

## Acknowledgements

We thank Kristina Grund, Kleintierpraxis Auringen, Germany, for veterinary care, and Ursula Wollscheid, Inge Lischewski, Lea Arndt and Anna Linck for excellent technical assistance. We also thank Christian Decker and Sebastian Görges for bioinformatic support.

## Supplementary

**Supplementary Tab. S1.** Available genome data used for comparison.

| Species | Accession Number |
| --- | --- |
| German Shepherd (*Canis lupus familiaris*) | GCA_011100685.1 |
| Labrador Retriever (*Canis lupus familiaris*) | GCA_012044875.1 |
| Basenji (*Canis lupus familiaris*) | GCA_013276365.2 |
| Cairn Terrier (*Canis lupus familiaris*) | GCA_031010295.1 |
| Bernese Mountain Dog (*Canis lupus familiaris*) | GCA_031010765.1 |
| Boxer (*Canis lupus familiaris*) | GCF_000002285.5 |
| Cat (*Felis catus*) | GCF_018350175.1 |
| Dingo (*Canis lupus dingo*) | GCF_003254725.2 |
| Mouse (*Mus musculus*) | GCF_000001635.27 |
| Human (*Homo sapiens*) | GCF_009914755.1 |

**Supplementary Tab. S2.** Repeat content of the Whippet genome assembly. Class, class of the repetitive regions. Count, number of occurrences of the repetitive region. bpMasked, number of base pairs masked; %masked, percentage of base pairs masked. LINE, Long Interspersed Nuclear Elements (including retroposons); LTR, Long Terminal Repeat elements (including retroposons); SINE, Short Interspersed Nuclear Elements; RC, Rolling Circle. In total, 902,477,158 bp were masked, corresponding to 36.5 % of the genome.

| Class | Count | bpMasked | %masked |
| --- | --- | --- | --- |
| SINEs | 478,438 | 65,159,195 | 2.64 |
| LINEs | 1,796,370 | 502,326,946 | 20.32 |
| LTR | 354,096 | 90,945,042 | 3.68 |
| DNA transposons | 300,163 | 49,080,228 | 1.99 |
| Rolling-circles | 1,568 | 78,131 | 0.00 |
| Unclassified | 525739 | 132,765,950 | 5.37 |
| small RNA | 61,933 | 5,023,647 | 0.2 |
| Satellites | 14,007 | 2,268,618 | 0.09 |
| Simple repeats | 870,502 | 48,791,971 | 1.97 |
| Low complexity | 109,450 | 6,037,430 | 0.24 |

**Supplementary Fig. S3.**
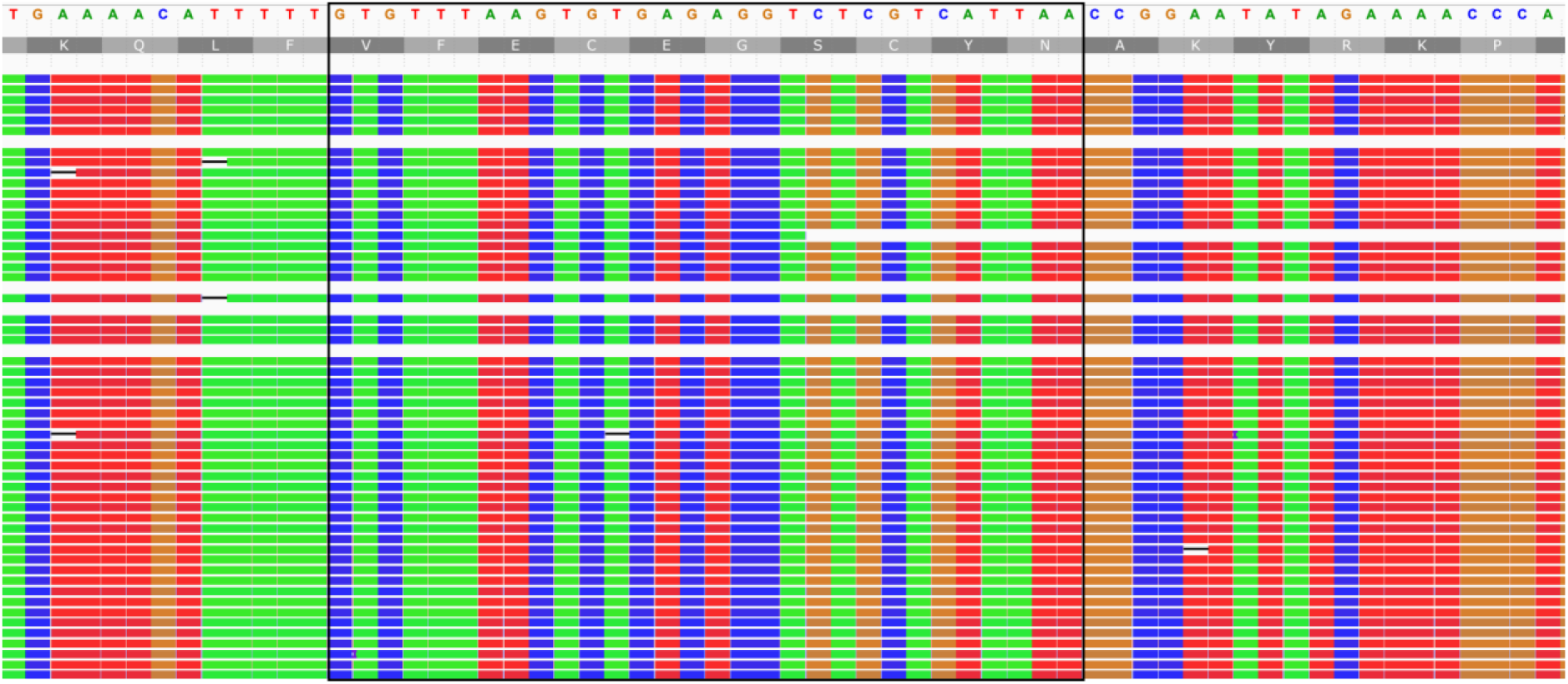
Read alignment of the myostatin gene, *MSTN*, in the sequenced Whippet genome. The black box indicates the *MSTN* sequence of interest, as used by Mosher *et al*. (2007).

## Notes

### Competing Interest Statement

The authors have declared no competing interest.

